# BRIDGE: A Coarse-Grained Architecture to Embed Protein-Protein Interactions for Therapeutic Applications

**DOI:** 10.1101/2025.04.11.648307

**Authors:** Pranav M. Khade, Andrew M. Watkins

## Abstract

In this work, we present Biophysical Representation of Interfaces via Delaunay-based Graph Embeddings (BRIDGE), a coarse-grained graph neural network that captures embeddings containing meaningful information about protein-protein interactions. The BRIDGE model takes as input graphs defined by the Delaunay tesselation of the individual chains and is pre-trained to predict the Delaunay adjacency at the protein-protein interface. The model achieves state-of-the-art performance in this task. The biophysical information captured by the BRIDGE embedding layer due to this pre-training task can further be used for downstream tasks, including for therapeutically relevant property prediction. We demonstrate the use of these embeddings by training models to predict antibody-antigen (Ab-Ag) affinity and antibody viscosity. These predictors, which require only the structures of the individual molecules in isolation, are especially well suited to large molecule therapeutic applications where complex structures are rarely determined until the very latest stages of a project.

## 1 Introduction

Biophysical interactions between binding proteins are critical for many biological processes. Capturing and characterizing these interactions can be valuable for many therapeutic and non-therapeutic areas involving protein design.

Although there are known biophysical interactions that warrant strong binding between the PPIs such as the presence of hydrophobic clusters, salt bridges, hydrogen bonds, etc. [1], simple models of protein-protein binding affinity, relying only on those well understood interactions cannot reliably achieve chemical accuracy, likely because influences from the conformational ensemble and interactions poorly captured by this classical decomposition are sufficient limitations. Since the precise, classical approach remains limited, a heuristic approach towards capturing these complex biophysical interactions can offer important benefits. In this study, we create a coarse-grained graph neural network called BRIDGE, with the goal of capturing biophysical properties, particularly those most relevant to protein-protein interactions, using a simple pretraining task of interchain residue contact prediction to generate embeddings.

There are several protein sequence [2, 3] and many structure-based [4, 5] protein interface prediction models that are built specifically for PPI prediction. Because structure prediction of single domains is now relatively reliable, we elect to build a structure-based model using *apo* structures, and to use a coarse-grained model to avoid excessive sensitivity to structural details that may be unreliably predicted. We leverage the DIPS-Plus [6] database, a comprehensive collection of experimentally derived protein structures, ensuring the robustness and reliability of our approach. Employing Delaunay tessellation [7] as the basis for graph construction, we minimize noise and spurious connections [8] between amino acid residues, enabling the generation of embeddings that reflect genuine biophysical interactions. By filtering extraneous spatial relationships, this strategy enhances the structural interpretability and predictive performance of the resulting embeddings.

The BRIDGE, a coarse-grained graph neural network (GNN) architecture (Figure 1), incorporates advanced layers to capture both local and global structural features of proteins. The model achieves an F1 score of 0.84 in interchain edge prediction. This performance underscores the efficacy of our method in learning representations that encapsulate meaningful biochemical and spatial properties. Furthermore, we demonstrate the use of these embeddings in therapeutic space by applying them to Antibody (Ab) - Antigen (Ag) affinity and antibody viscosity prediction and outperforming the state-of-the-art models in the case of viscosity prediction and impressive performance despite fewer training data points in the case of affinity prediction.

**Fig. 1.**
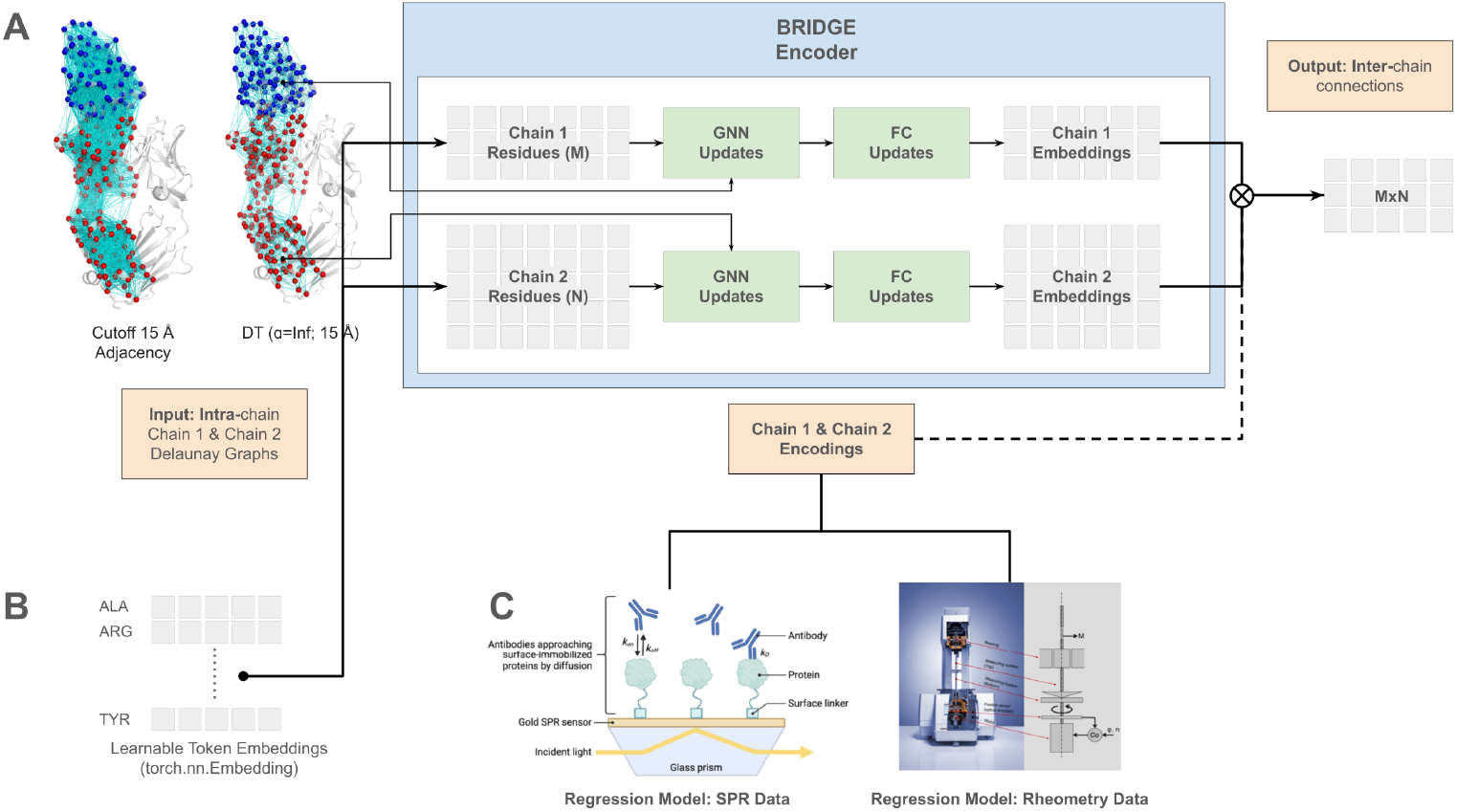
BRIDGE architecture uses learnable node embeddings and Delaunay adjacency as an input. The primary task (output) for the model training is to predict the interacting residues at the interface (Delaunay edges between the two graphs of two chains; ⊗ symbol represents the outer product.). While learning this task, the model embeddings capture critical biophysical information about protein-protein interfaces that can be repurposed in therapeutic space. (A) BRIDGE uses Delaunay tessellations as adjacency with cutoff 15 Å. This significantly reduces the number of ‘spurious’ edges and makes training more efficient. For example, the antibody-antigen structure (PDB ID: 6O39) has 7,959 residue pairs within 15 Å in total, only 2,410 of which are in the Delaunay tesselation, a 69.7 % reduction in the number of edges. (B) The sequence of each amino acid chain in the protein complex is embedded; the details of the model hyperparameters can be found in Table S1 (C) The BRIDGE embeddings can be repurposed for various therapeutic applications. To demonstrate this, we have trained regression models to predict the antibody-antigen interaction strength (SPR data) and also the viscosity of high-concentration antibody solutions (rheometry).

By combining experimental data with geometric graph-based methods, our approach highlights the potential of embeddings derived from PPI graphs to serve as powerful tools for diverse applications in therapeutic space.

## 2 Results

### 2.1 BRIDGE PPI prediction performance

Model performance on the edge prediction pretraining task was assessed using the Validation F1 score, with an observed score of 0.84. More metrics are listed in Table 1. The SpatialPPI 2.0 [5] model, which similarly predicts the interface contact map, achieves state of the art performance on the PINDER dataset [9]. Since PINDER includes a large number of AlphaFold2 examples [10, 11], we decided to use the DIPS-Plus [6] database which includes only experimental structures. SpatialPPI 2.0 is predicting an all-residue contact map rather than only Delaunay contacts, and since the training set includes some lower-confidence structures, so we do not see the results as precisely comparable: perhaps Delaunay contacts are somewhat harder to predict correctly, while correct prediction on AlphaFold2 models is more challenging. We include the SpatialPPI 2.0 results in 1 as the closest point of reference we have available, but between the somewhat different evaluations and the use of this task only to obtain meaningful embeddings, we are mostly interested in suggesting that we have successfully learned something significant about interfaces.

**Table 1.** Validation set metrics for the BRIDGE encoder trained on DIPS-Plus dataset with comparable results for SpatialPPI 2.0 on PINDER dataset.

| Score | BRIDGE (DIPS-Plus) | SpatialPPI 2.0 (PINDER) |
| --- | --- | --- |
| F1 | 0.84 | 0.732 |
| Accuracy | 0.85 |  |
| Precision | 0.919 | 0.698 |
| Recall | 0.775 | 0.769 |
| Average Precision | 0.825 | 0.783 |
| AUROC | 0.854 |  |

### 2.2 BRIDGE-Affinity Prediction

To test the embeddings in the therapeutic domain, we conducted a UMAP analysis of the independent dataset (SAbDab). The main objective of the analysis was to test if the model captures biophysical knowledge from the link prediction task and if that can be used for tasks such as affinity prediction. We conducted dimensionality reduction using UMAP before building a regression model to check if embeddings form clusters that indicate biophysical information in the embeddings. Figure 2 A shows that we observe fairly separated clusters of low and high-affinity data points. However, there are multiple cluster centers instead of two, since the embeddings likely capture substantial structural signal unrelated to affinity. The hyperparameters for the UMAP analysis are listed in Table S4.

**Fig. 2.**
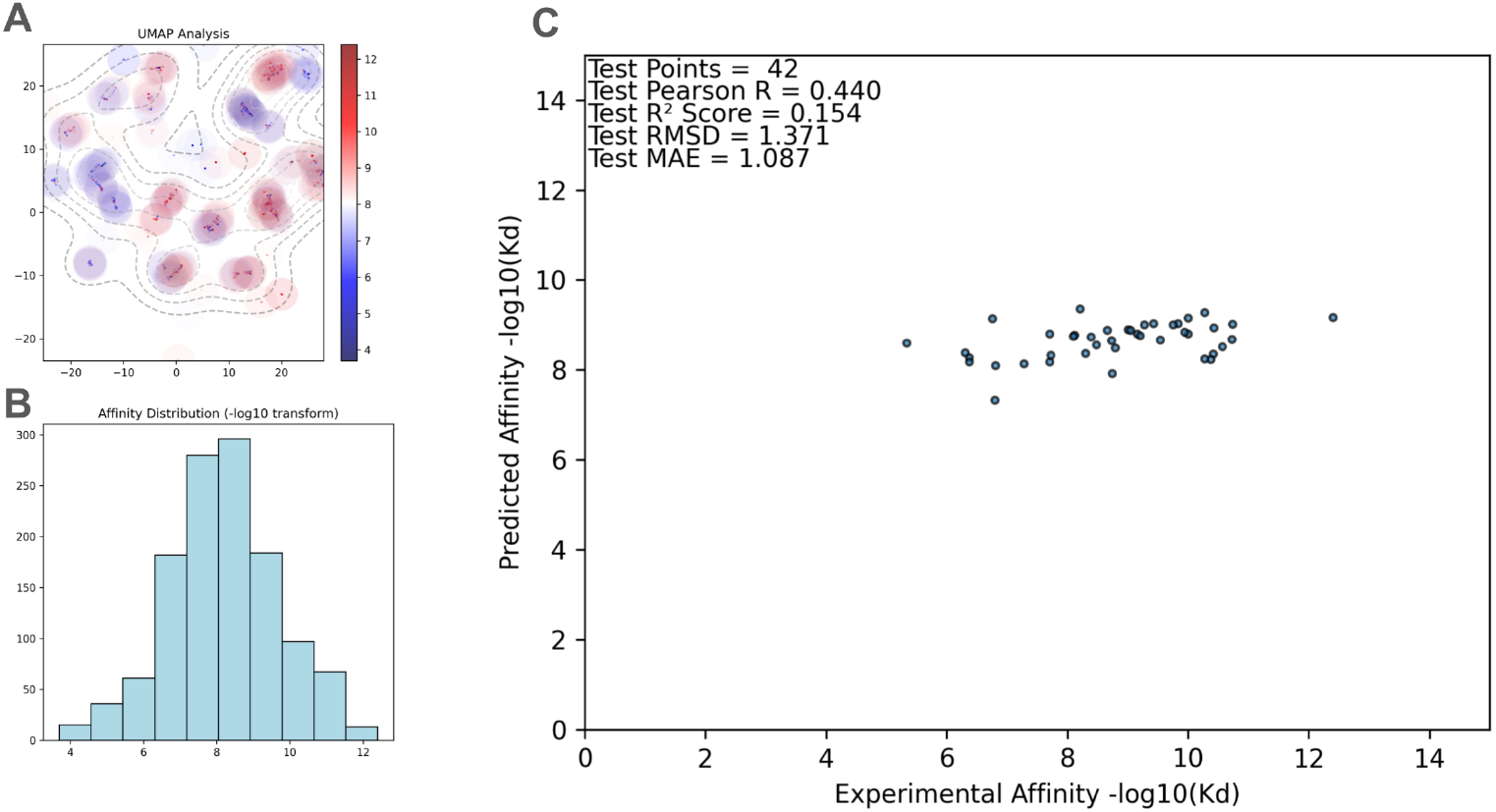
SAbDab SPR measurement data regression experiment (A) Dimensionality reduced embeddings (UMAP) obtained after parsing Ab & Ag graphs through the BRIDGE Model. There are some clusters of high and low affinity but many without a clear boundary and consensus (unlike viscosity analysis) which demonstrates the complex and hard-to-categorize nature of biophysical phenomena involved in protein-protein binding. (B) Affinity values have a lognormal distribution which although makes it difficult to model the extreme values, it also helps the model to understand subtle differences that cause changes in affinity around the mean value. (C) BRIDGE-Affinity performs modestly on the held-out Pierce lab dataset.

Since the UMAP analysis suggests BRIDGE embeddings may capture some information about molecular interactions, we also built a regression model that uses embeddings to predict the affinity of Ab-Ag complexes. Embeddings of the two chains are used to calculate the outer product and it is flattened to be used as an input to a fully connected feed-forward neural network. The regression model (Figure 2 C) performs better on the held-out Pierce lab dataset than any model but DG-Affinity [12], which had access to an additional non-public training dataset (Table 2).

**Table 2.** Validation metrics, as well as held-out test set metrics, for compared models. The performance of other top-performing methods is directly taken from the DG-Affinity paper. While DG-Affinity was trained on single-domain antibodies and a privately held dataset called “PaddlePaddle,” BRIDGE-Affinity was trained and validated on 938 de-duplicated antibodies from SAbDab.

| Method | Validation |  | Pierce Lab Data |  |
| --- | --- | --- | --- | --- |
| | Pearson R | $R^2$ Score | Pearson R | $R^2$ Score |
| BRIDGE-Affinity | **0.697** | **0.364** | 0.44 | 0.154 |
| DG-Affinity* | 0.602 | 0.353 | **0.656** | **0.367** |
| ZRANK | 0.3252 | -1663.9615 | 0.368 | -2257.33 |
| AP PISA | 0.3145 | -30.4824 | 0.414 | -39.67 |
| AP DCOMPLEX | 0.3023 | -5.1368 | 0.172 | -4.843 |
| FIREDOCK AB | 0.2802 | -856.9465 | 0.345 | -1059.773 |
| ELE | 0.2753 | -33.3912 | 0.285 | -82.87 |
| AP T2 | 0.2716 | -672.8094 | 0.459 | -678.868 |
| PYDOCK TOT | 0.2559 | -160.5253 | 0.305 | -167.735 |

### 2.3 BRIDGE Viscosity Prediction

We have used two different strategies for the viscosity prediction using the same embeddings, data and architecture but different training strategies. One strategy is a 75:25 split and another with Leave One Out Cross-Validation (LOOCV). Rai et. al. [13] have developed and highlighted 3 independent models for these datasets with data augmentation and cross-validation. However, one of the model used both datasets (Ab21 and PDGF38) using the LOOCV strategy has a Spearman R score of 0.71 which is less than our model that uses the LOOCV strategy the representative model performance of the regression model is highlighted in Figure 3 C. However, detailed information along with comparisons with the Sharma et. al. model [14] and SCM [15] from the Rai et. al. paper are available in Table 3. It is important to note that for the BRIDGE Viscosity (75:25 split) model, we have not restricted it to predicting only low-viscosity points from Ab21, unlike other methods to truly gauge the model’s ability to predict viscosity in a wide range as well as with fewer data points. Furthermore, we have not used the docked form of the Abs which demonstrates that we do not need to obtain multimeric structures for pairwise analysis and docking step can be completely skipped as BRIDGE is trained on Delaunay edge prediction task that works as a meta-docking in graph space itself.

**Table 3.** Viscosity performance metrics for models. All the models except BRIDGE-Viscosity (75:25 split) mentioned in this table are trained on PDGF38 and Ab21 datasets with LOOCV strategy and tested only on Ab21. The BRIDGE-Viscosity model with (75:25) split has the most significant result even though the BRIDGE LOOCV model has better metrics since LOOCV uses more data and is more prone to overfitting.

| Method | Spearman R | $R^2$ Score |
| --- | --- | --- |
| BRIDGE-Viscosity (75:25 split) | 0.781 | 0.687 |
| BRIDGE-Viscosity (LOOCV) | 0.978 | 0.690 |
| Sharma | 0.55 | 0.48 |
| SCM | 0.49 | 0.40 |
| PfAbNet (LOOCV) | 0.71 | 0.79 |

**Fig. 3.**
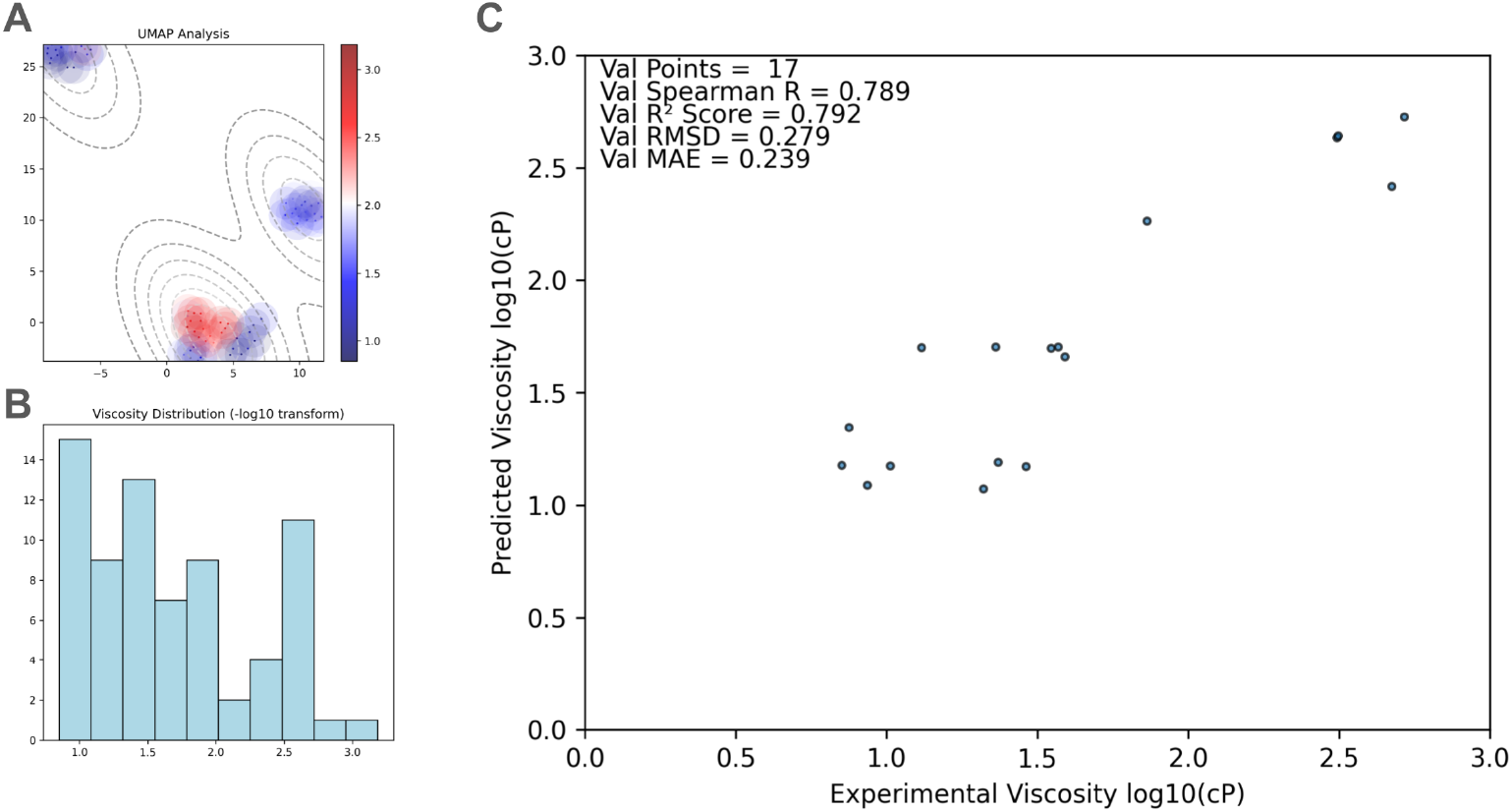
Regression modeling of high-concentration antibody viscosity. (A) Dimensionality reduced embeddings obtained after parsing the same Delaunay graph (obtained from a predicted Fv structure) for both chain inputs to the BRIDGE model: implicitly, we model Fv/Fv interactions and their contribution to viscosity. We see a clear distinction between the viscous and nonviscous clusters, with few exceptions. (B) Viscosity values are distributed more evenly than affinity values. (C) Performance of BRIDGE-Viscosity (75:25 split) model; We have put a wide range of viscosity examples in training as well as the validation sets instead of focusing on the Ab21 dataset containing only low viscosity values. This single model outperforms state-of-the-art models with the Spearman Correlation coefficient as a metric without any data augmentation strategies.

Upon investigating the reduced dimension space of the embeddings obtained using UMAP (Figure 3 A) we see distinct clusters of low and high-affinity values which explain the good performance of the regression model. Furthermore, the viscosity values are evenly spread (Figure 3 B) which means the clusters are not formed because of a sheer number of similar values and are more likely to be formed because of the interaction surface biophysical similarity. The hyperparameters for the UMAP analysis are listed in Table S4.

## 3 Methods

### 3.1 Datasets and Delaunay Tessellation Graph Construction

#### DIPS-Plus database

A total of 42,112 protein chain pairs were selected from DIPS-Plus [6], ensuring diverse structural and interaction characteristics by using the default train-validation splits. Each chain in the pair was a coarse-grained model represented by its alpha carbon (C*α*) atoms to create a simplified yet accurate model of spatial relationships. Delaunay tessellations [7] were obtained from each protein chain’s C*α* atom coordinates. This approach creates non-overlapping tetrahedra (converted to four edges) that link nearby residues while omitting irrelevant or intervening connections, effectively capturing local packing and interaction environments within each protein chain [8] (Figure 1) A. Also, a reduction in the number of edges means more efficient GNN model training [16]. We cut off the edges between the C*α* atoms that had a distance of more than 15 Å; We have used this adjacency definition for all the models in this paper.

#### Antibody-Antigen Affinity Dataset

We used a subset of the SAbDab [17] (1231 data points) database that contained Surface Plasmon Resonance (SPR) affinity values associated with it. We further filtered the database by excluding entries with synthetic amino acids (To avoid creating extra tokens) and missing chains.

#### Antibody Viscosity Dataset

We have used the same data from Rai et al. [13]: a dataset containing experimental viscosity measurements of 27 FDA-approved therapeutic antibodies (Ab21 set). (Rai et. al. used IgG1 only filter resulting in 21 entries that we did not filter out for the 75:25 split model but filtered out for the LOOCV strategy), 38 anti-PDGF antibody variants (PDGF38 set) developed in a Pfizer-internal program and an additional 7 antibodies (Basiliximab, Natalizumab, Tremelimumab, Ipilimumab, Atezolizumab, Ganitumab, Vesencumab) with available sequences.

### 3.2 BRIDGE Architecture (PPI Prediction)

#### Model Overview

Two independent stacks of GNN layers and fully connected layers are used to generate chain-specific embeddings. This model was trained with the task of predicting inter-chain protein-protein interactions given two Delaunay graphs of chains.

#### Node Features and Edge Definition

Each residue node was initialized with learnable embeddings. There are a total of 27 different tokens that include 20 standard amino acids, 6 ambiguous (UNK, ASX, GLX, XAA, XLE), non-standard amino acids (SEC, PYL), and 1 gap token.

#### Embedding Generation

Separate embeddings were generated for each chain using a series of graph convolution layers including GATv2Conv [18], GCNConv [19] and fully connected layers the architecture is shown in Figure S1, designed to capture both local and global structural information within each protein. Layer normalization [20] and dropout were applied after each layer to improve model generalization and mitigate overfitting.

#### Embedding Combination

Following individual chain processing, the embeddings from each chain were used to predict the interchain connections using positive examples and negative sampling with an equal number of negative labels. For the probability of a connection, embedding for individual amino acids from different chains is used to calculate the dot product which is then parsed to the Sigmoid layer for binary label prediction. The schema for this process is in Figure S1.

#### Training and Evaluation

The model was trained with binary cross-entropy loss to optimize interaction prediction.

### 3.3 BRIDGE Affinity Prediction

We used the trained BRIDGE model to obtain embeddings for Ab-Ag structures by parsing the Delaunay graphs and we use the output embeddings to calculate the outer product of two embedding vectors and flatten it to get a pair representation. Firstly, we removed entries with 42 Ab-Ag antibody PDB IDs overlapping with the Pierce Lab dataset [21] from the training and validation set, which was later used for testing. We then split the remaining 1172 structures into training (1060 entries) and validation (112 entries) so that no PDB ID and affinity combination present in the training dataset is found in the validation dataset. We call this process ‘deduplication’. We then use the regression architecture shown in Figure S2 to fit the Ab-Ag affinity with Huber loss with weights according to the PDB ID frequency (non reduced loss function is directly multiplied by frequency of PDB ID) and trained models with random splits five times to get average performance.

### 3.4 BRIDGE Viscosity Prediction

To obtain embeddings for this model, we used ABodyBuilder2 [22] on the antibody sequences from the viscosity database to obtain Fab variable (fv) domain structures. We then parse the same antibody Delaunay Graph twice to the BRIDGE model to get self-binding embeddings as viscosity is known to be the result of mainly antibody self-interactions. We calculate the outer product of the embeddings and form the same pair representation as the BRIDGE-Affinity. We have developed a single model to predict the viscosity with identical architecture to that of the BRIDGE-affinity model (Figure S2) and used 75:25 train: validation split. We also ensure representation from each dataset (Ab21, Ab8 and PDGF38) in training and validation, we split all datasets in the same proportion. We have another training strategy using LOOCV for the Ab21 and PDGF38 combined dataset to make direct comparisons with the Rai et. al. study. For the BRIDGE LOOCV model, we use five seeds (Table S3) and select the top three checkpoints with the lowest loss (Total 5*3 = 15 models for each example) to predict viscosity and take the median of the predictions to get a single value for an example. We also didn’t log-transform the viscosity values, unlike the 75:25 split model.

### 3.5 Experimental Setup and Hyperparameters

#### Training Parameters

The BRIDGE model was trained using hyperparameters mentioned in Table S1. Hyperparameters for the BRIDGE-Affinity and BRIDGE-Viscosity models are mentioned in Table S2 and Table S3 respectively.

#### Computational Resources

The model is trained on NVIDIA Quadro P6000 GPU with PyTorch, PyTorch-Lightning, and PyTorch Geometric [23].

## 4 Discussion

Pretraining a Delaunay graph encoder on interface edge prediction yields biophysically meaningful descriptors useful for downstream tasks. Beyond immediate use cases where experimental data is ordinarily limiting (such as antibody-antigen complex structures necessary for accurate affinity estimation), this study suggests that antibody-antibody self-interaction can be distilled in much the same way to explain some variation in viscosity. By using the Delaunay tessellation, the BRIDGE model can predict fewer edges and can use far fewer parameters (*<*1M, in contrast to language model embeddings like ESM 3 [24], which has up to 98 billion trainable parameters). Biological priors distilling structural insight can use computational resources more efficiently.

These embeddings may prove useful for further biological tasks, such as binder generation or protein interaction prediction on large sequence databases like Negatome.

## 5 Conclusion

Conclusions In this study, we introduced a coarse-grained BRIDGE architecture that effectively captures biophysical embeddings from PPIs. By leveraging Delaunay tessellation for graph construction and training on experimentally derived structural data, our model not only achieves state-of-the-art performance in interchain interaction prediction but also encodes meaningful biophysical properties in its learned embeddings. These embeddings were further demonstrated to be valuable for downstream tasks, such as affinity prediction, and viscosity prediction in therapeutic applications. Our approach addresses the challenges associated with limited structural data in the therapeutic protein domain by utilizing a broader set of non-therapeutic protein structures for training. The BRIDGE model’s efficiency, with only 847K trainable parameters, underscores the potential of targeted, computationally efficient architectures in extracting meaningful representations from protein interfaces. Beyond the demonstrated applications, the extracted embeddings offer a foundation for future explorations, including generative modeling and broader applications. Additionally, the modular nature of our framework allows for easy extensions, such as incorporating non-amino acid entities like small molecules and nucleic acids, further broadening its utility. Overall, this work highlights the power of structure-based graph learning for capturing and leveraging biophysical principles in protein interactions, paving the way for novel insights in structural biology and drug discovery.

## Acknowledgements

We sincerely thank Dr. Sai Pooja Mahajan for their invaluable advice and insightful suggestions, which greatly contributed to the development and refinement of this research.

## Disclosure

All authors are employees of Genentech and some may be shareholders of Roche.

## Data Availability Statement

Code to reproduce all the experiments and to train the BRIDGE model is available at the following link: https://github.com/prescient-design/BRIDGE

## Notes

https://github.com/prescient-design/BRIDGE

