## Supplementary Material for "BRIDGE: A Coarse-Grained Architecture to Embed Protein-Protein Interactions for Therapeutic Applications"

**Learnable Node Embeddings.** We analyzed the pre-network amino acid node embeddings that have a total of 27 tokens (not to be confused with protein pair level embeddings) for the model with the best F1 score. We analyze these embeddings to search for biologically relevant patterns. Note that these embeddings are not standalone but still provide valuable insights. Embeddings were analyzed with cosine similarity. We also clustered the embeddings using UMAP (Fig S3). For simplicity, we present results only on 20 standard amino acids here.

While analyzing the amino acid embeddings, we want to highlight that we do not expect results to be similar to BLOSUM Biophysical properties such as those categorized in Fig S3 B, note that these embeddings are pre-network and a lot of contextual information will be missing from them since it will be captured in the multihead GATv2Conv layer attention that we are using in our architecture.

The analysis reveals distinct clustering patterns for specific amino acid types in both the heatmap and scatterplot representations. Nonpolar, aliphatic amino acids, such as LEU and MET, form tight clusters, reflecting their similar properties in protein structures. Positively charged amino acids, including ARG and LYS are positioned close to each other, highlighting their shared electrostatic characteristics. Similarly, aromatic amino acids, such as PHE and TYR, exhibit proximity in both visualizations, emphasizing the structural and chemical similarities of their R-groups.

### S1 Supplementary Tables

| Hyperparameter | Value | Description |
|---|---|---|
| Learning Rate | 0.001 | ExponentialLR is used with $\gamma$: 0.98. |
| Epoch | 289 | |
| Batch Size (T/V) | 64/32 | Training/ Validation |
| Seed | 42 | Reproducibility |
| Amino Acid Embedding Vector Length | 150 | |
| Hidden Dimension | 400 | |
| Probability threshold for connection | 0.75 | |
| GATv2Conv heads | 3 | |

**Table S1** BRIDGE hyperparameters used to build the PPI prediction model with 847 K parameters.

| Hyperparameter | Value | Description |
|---|---|---|
| Learning Rate | 0.00008 | ExponentialLR is used with $\gamma$: 0.995. |
| Epoch | 103 | |
| Batch Size (T/V) | 32/16 | Training/ Validation |
| Seed | 42 | Reproducibility |
| Hidden Dimension | 50 | |

**Table S2** BRIDGE-Affinity hyperparameters used to build the affinity regression model with 70M parameters. This model has high parameters because of the flattening & padding we have to do for the embedding dot product matrix.

| Hyperparameter | Value | Description |
|---|---|---|
| Learning Rate | 0.01 | ExponentialLR is used with $\gamma$: 0.99. |
| Epoch | 215 | |
| Batch Size (T/V) | 55/17 | Training/ Validation |
| Seed | 42 | Reproducibility |
| Hidden Dimension | 50 | |
| LOOCV additional parameters | | |
| Seeds | 239, 1223, 2342, 5290, 234 | Reproducibility |

**Table S3** BRIDGE-Viscosity hyperparameters used to build the affinity regression model with 2.7M parameters.

| Pre-BRIDGE-Affinity UMAP Analysis | | |
|---|---|---|
| Hyperparameter | Value | Description |
| Components | 2 | Reduction to 2D |
| Neighbors | 10 | |
| Minimum Distance | 0 | |
| Metric | Canberra | |
| Initialization | Random | |
| Epochs | 500 | |
| Negative Sample Rate | 20 | |
| Random State | 42 | Reproducibility |
| 'y' Values Provided | True | |
| Pre-BRIDGE-Viscosity UMAP Analysis | | |
| Components | 2 | Reduction to 2D |
| Neighbors | 15 | |
| Minimum Distance | 0 | |
| Metric | Canberra | |
| Initialization | Random | |
| Epochs | 500 | |
| Negative Sample Rate | 20 | |
| Random State | 42 | Reproducibility |
| 'y' Values Provided | False | |

**Table S4** UMAP Hyperparameters

### S2 Supplementary Figures

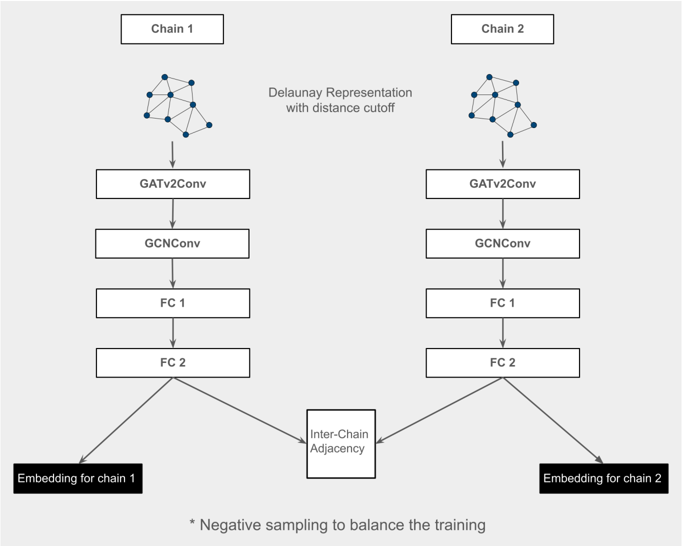


**Fig. S1** Fig. BRIDGE PPI prediction model schema. The model has only 847K trainable parameters.

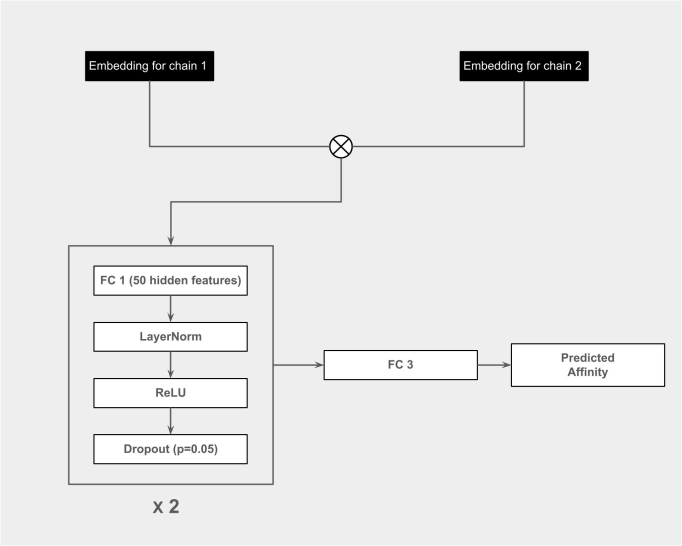


**Fig. S2** Fig. BRIDGE-Affinity prediction model schema. The ⊗ symbol represents the outer product.

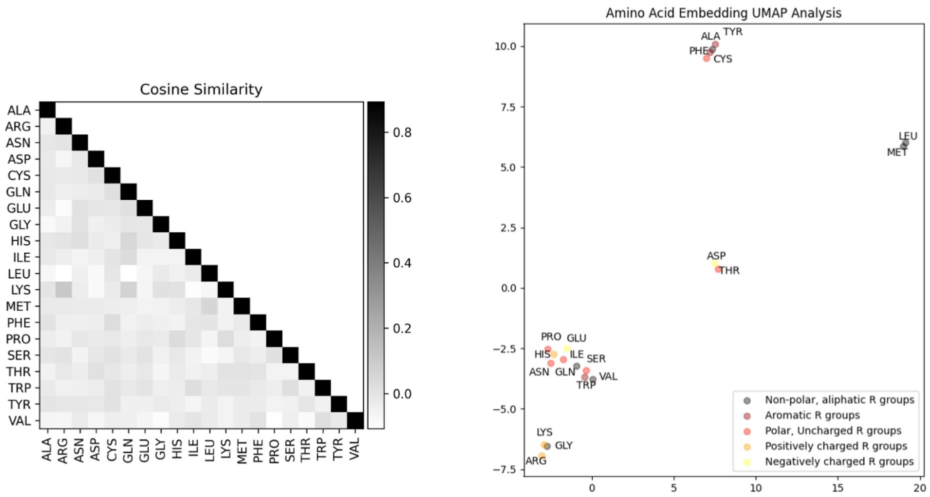


(A) (B)

**Fig. S3** Learned amino acid embedding (pre-GAT attention) analysis shows learned biophysical similarity. (A) Pairwise cosine similarity between the embeddings. (B) UMAP is initialized with principle component analysis (PCA) and run for 50 epochs also, cosine similarity is used as a metric; the value of two is selected for the nearest neighbors parameter.

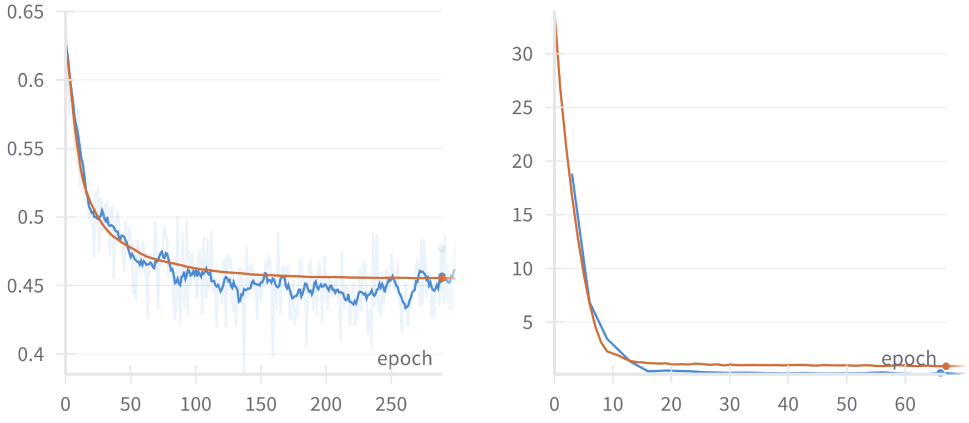


(A) (B)

**Fig. S4** Loss function over epochs for (A) PPI prediction model (BCE loss) with final model training loss (blue) 0.457 and validation loss (red) 0.455. The training loss is smoothed with the running average. (B) Affinity regression model (MSE loss) with final model training loss (blue) 0.26 and validation loss (red) 0.88.
